# Group social dynamics in a semi-natural setup reveal an adaptive value for aggression in male mice

**DOI:** 10.1101/2024.04.25.591070

**Authors:** Sergey Anpilov, Yair Shemesh, Asaf Benjamin, Tommaso Biagini, Yehezkel Sztainberg, Alon Richter-Levin, Oren Forkosh, Alon Chen

**Affiliations:** Department of Brain Sciences, Weizmann Institute of Science; Rehovot, 7610001 Israel; Department of Molecular Neuroscience, Weizmann Institute of Science; Rehovot, 7610001 Israel; Department of Cognition and Brain Sciences, The Hebrew University; Jerusalem, 9190501 Israel

**Keywords:** Aggression, Ethology, Hierarchy, Oxytocin, Semi-Natural

## Abstract

**Background:** Maladaptive aggression in humans is associated with several psychiatric conditions and lacks effective treatment. Nevertheless, aggression constitutes an essential behavior throughout the animal kingdom as long as it is tightly regulated. Studying how social dominance hierarchies (SDH) regulate aggression and access to resources in an enriched environment (EE) can narrow the translational gap between aggression in animal models and humans normal and pathological behavior.

**Methods:** The ‘social box’ (SB) is a semi-natural setup for automatic and prolonged monitoring of mouse group dynamics. We utilized the SB to decipher complex tradeoffs between aggression, social avoidance, resource allocation, and dominance in two mouse models of increased aggression: (i) a model of early exposure to EE and (ii) a model of oxytocin receptor deficiency (OxtR^-/-^). While EE increases aggression as an adaptive response to external stimuli, hyper-aggression in OxtR^-/-^ mice is accompanied by marked abnormalities in social behavior.

**Results:** EE groups exhibited significant social avoidance, and an increased proportion of their encounters developed into aggressive interactions, resulting in lower levels of exploratory activity and overall aggression. The hierarchy in EE was more stable than in control groups, and dominance was correlated with access to resources. In OxtR^-/-^ groups, mice engaged in excessive social encounters and aggressive chasing, accompanied by increased overall activity. In OxtR^-/-^ groups, dominance hierarchies existed but were not correlated with access to resources.

**Conclusion:** Measuring aggression and social dominance hierarchies in a semi-natural setup reveals the adaptive value of aggression in EE and OxtR^-/-^ mice, respectively. This approach can enhance translational research of pathological aggression.

## Introduction

Aggression is an easy and intuitive behavior to detect in humans and other animals, including mice in the laboratory. However, assessing whether aggressive behavior is beneficial or detrimental poses a complex challenge (1). In humans, aggression can be either instrumental or hostile (2); whereas *instrumental aggression* is a calculated effort to obtain desired objectives, hostile aggression is a spontaneous, impulsive act aiming to harm another person (3). Theoretical models further delineate pathological aggression as an extreme and unreasonable act, relying on multiple criteria such as personality and context (4). In contrast to human studies, in ethological research, pathological aggression is not applied even to deadly aggressive acts, which are observed as legitimate survival-oriented measures (5). Therefore, maladaptive human aggression currently lacks an appropriate animal model, potentially hampering the translation of research findings to human applications.

Most preclinical studies measure aggression in the ‘resident-intruder’ test (RI), quantifying the readiness of a male mouse to defend its cage (the ‘territory’) from an unfamiliar intruder (6). The RI test has induced significant advances in describing the central aggression network (7). Nevertheless, from an ethological perspective, short dyadic paradigms that model aggression in a relatively specific context might have limited translational value (8).

Mice are social and territorial species with high fertility rates, dispersal, and competition. As opportunists, mice adapt to diverse habitats, and commensal mice that live near human dwellings, where food resources are patched and concentrated, display more fierce territorial conflicts than feral mice (9). In agreement with that, in laboratory settings, aggression increases with the enrichment of the setup (10). For instance, there is an increase in inter-male aggression and the frequency of injuries from biting in more vast and crowded arenas (11). Similarly, biting injuries are more common in groups of males following prolonged post-weaning housing in an enriched environment (12). Though aggression during the competition for scarce resources such as food, mates, and shelter is beneficial, it is also risky and must be tightly controlled. Establishing social dominance hierarchies (SDH) minimizes aggression by setting a priority order of access to desirable resources, thereby avoiding unnecessary conflicts (13). Thus, dominance hierarchy could be set as a reference measure for delineating maladaptive aggression in preclinical studies in laboratory settings (14). SDH in mice is commonly tested using the ‘Tube Test’ (15) however, simultaneously measuring aggression and SDH in the same group of mice using the RI and Tube tests is not feasible.

We postulated that monitoring aggression and SDH in groups of mice in a semi-natural setup can bridge the translational gap and help elucidate mechanisms underlying pathological aggression (16). Our group developed a semi-natural setup called the ‘Social Box’ (SB) (17). In the SB paradigm, mice with different color-marked fur are uninterruptedly maintained in a large, complex arena and video-recorded over several days. Next, the videos are subjected to automatic tracking of the individual mice, enabling the extraction of multiple behavioral readouts, such as aggressive chasing and escaping (18–20). Consistent within-group imbalance of chase/escape behavior is further mathematically expressed to represent SDH (21).

Here we utilized our semi-natural setup to test both social and individual behavioral characteristics in two mouse models of increased aggression: i) aggression due to early and prolonged exposure to environmental enrichment (EE) and ii) aggression due to developmental deficiency of the oxytocin receptor (OxtR) (22,23). Early exposure to EE physiologically benefits the individual and encourages ingroup competition (11). Mal-function of the oxytocinergic system due to OxtR deficiency alters social behaviors, including aggression (24–26). We hypothesized that the EE-induced aggression would reflect an adaptation contributing to group coherence as measured by social interactions, hierarchy, and exploration in the SB. In contrast, since the hyper-aggressive phenotype of OxtR^-/-^ mice is not shaped by malfunctioning the oxytocin system and not by environmental needs, we hypothesized that this manipulation would impair group coherence in the SB.

## Methods and Materials

### Animals

The Institutional Animal Care and Use Committee of The Weizmann Institute of Science approved all experimental protocols. For the EE experiment, adult male ICR mice (Harlan Laboratories, Jerusalem, Israel) were used. For the OxtR^-/-^ experiment Mice, initially generated by Lee et al. (23) carrying the mutation on the genetic background of the 129/Sv line were crossed with ICR mice. Experimental mice were produced by crosses of heterozygous F1 parents. To form experimental groups of four male mice, two pairs of littermates obtained from a separate family were weaned into the same cages. Throughout both experiments, the animals were maintained in a temperature-controlled mouse facility (22°C ± 1) on a reverse 12 hr light–dark cycle. Food and water were given *ad libitum*. Notably, the mice in the two experiments differ in their genetic background (outbred in the EE experiment and mixed background in the OxtR^-/-^). Furthermore, parental OxtR haploinsufficiency might affect parameters such as parental care. Hence, we compared the treatment to its control within each experiment and avoided quantitative comparisons across experiments.

### Color markings of the fur, the social box setup, and mice tracking

As described in Shemesh et al. 2013 (17) (see Supplemental Information).

### Identification and classification of behavior

We defined a set of 22 a priori chosen behavioral readouts as described in Anpilov et al. 2020 (19), which included both individual and pairwise automatically detected behavioral readouts (see Supplemental Information)

### Quantification and statistical analysis

#### Multi-variate separation

calculated as Euclidean distance between the Cre+ and Cre-mice centroids in 21 dimensions of the automatically collected behavioral readouts. Statistical inference was based on comparison to a null distribution of Euclidean distances corresponding to 1×10^5^ randomly reshuffled partitions.

#### Principal component analysis (PCA)

was performed as described in Anpilov et al. 2020 (19) (see Supplemental Information).

#### The logistic regression model

To quantifies the relationship between the group types (EE vs. SC and OxtR^-/-^ vs. OxtR^+/+^) and the behavioral readouts measured, logistic regression models were fit with group type as the label and behavioral readouts as the features, using the Matlab (2021b) function ‘fitclinear’ with Lasso regularization and the default parameters. Leave-one-out cross-validation (CV) was used to estimate the performance of these models. Next, the value of the regularization parameter was chosen based on CV performance to avoid overfitting and assert that only truly predictive features will have significant coefficients, and the coefficients of these well-regularized models and their 95% confidence intervals were then used to examine the relationship between the variables. Importantly, to remove potential colinearity between the behavioral readouts, the behavioral data were first whitened using zero-phase component analysis (ZCA), and the models were fit to these whitened data.

#### David score calculation, Steepness of hierarchy, Despotism, and Directional consistency

as described in Karamihalev et al. 2020 (20) (see Supplemental Information).

#### Elo rating

By treating chasing and escaping as ‘wins’ and ‘losses’ the rank over time can be quantitatively represented by the Elo rating. Briefly, every participant is assigned the same initial rating. Upon each subsequent encounter, the ratings of the winner and the loser are symmetrically adjusted according to the outcome and proportionally to the difference between the current ratings (see Supplemental Information).

#### Permutation test

for inferential comparisons between two samples, the Monte Carlo permutation test of means (1×10^7^ permutations) was carried out,

All the above analyses were performed using MATLAB (R2019a, MathWorks, Natick, MA, USA).

## Results

### Exploratory and social activity is decreased following adolescence exposure to EE but increased in OxtR deficiency

#### Aggression in the SB due to early and prolonged exposure to EE

At the age of weaning (3 wks), mice were randomly distributed into two types of groups: standard conditions (SC) mice that were housed in groups of four in standard laboratory cages, and enriched environment (EE) mice that were housed in groups of 16 male mice in a spacious and complex cage, with a variety of objects such as shelters, tunnels, running wheels, and mouse nest boxes (27) (Figure 1A). Six weeks later, 4 out of 16 mice in each group were randomly chosen to constitute the treatment groups (EE), and the fur of each of the four mice was colored differently. These four mice were returned to the enclosure for an additional week, while the rest of the subjects were omitted from the experiment. Then, at the age of 10 weeks, the group was introduced into the SB for uninterrupted tracking for 12 hours each day during the dark phase over 4 consecutive days. As a control, we used eight groups of four male mice housed in a home cage (‘standard condition’; SC) until they were introduced to the SB at 10 weeks (Figure 1A).

**Figure 1.**
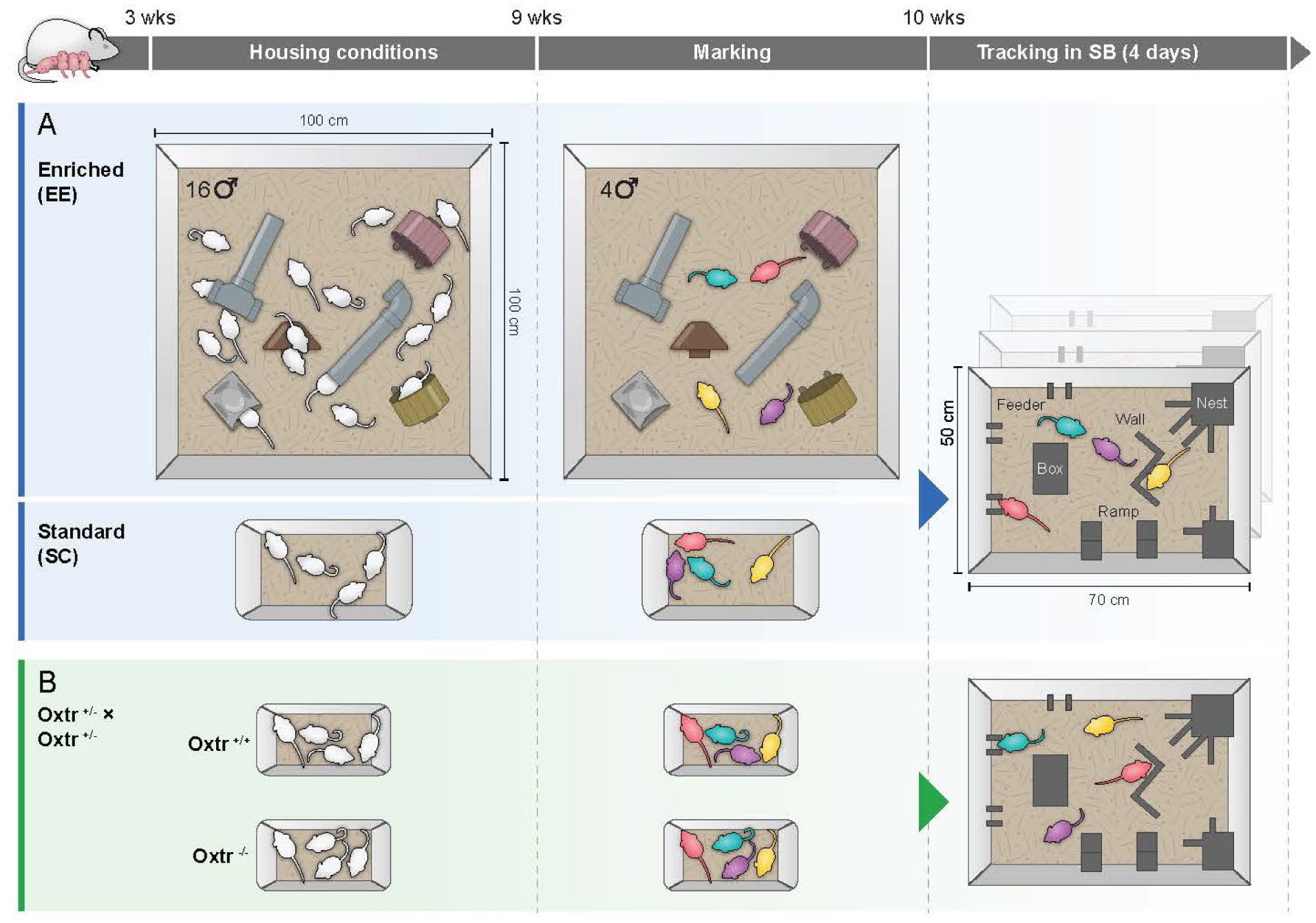
Experimental design. **(A)**. Enriched environment (EE) groups of 16 male mice were kept in the EE cages during adolescence; four of them were marked, regrouped, and reintroduced to the EE cages for a week and transferred to the SB at the age of 10 weeks. **(B)** Oxytocin receptor deficient mice (OxtR^-/-^) were kept in groups of 4 males in a standard home-cage until transfer to the SB at the age of 10 wks. The SB arena was divided into discrete regions of interest-open area, Z wall, water, feeders, large nest, block, and small nest.

#### Aggression in the SB due to OxtR deficiency

Eleven groups of OxtR^-/-^ (treatment) and eight groups of OxtR^+/+^ (control) weanlings were assembled from heterozygous breeding (OxtR^-/+^ × OxtR^-/+^) of mixed background. Each group consisted of four males kept in a standard home cage for 7 wks. Thereafter, at the age of 10 weeks, the groups were introduced to the SB for 4 days (Figure 1B).

First, we inspected ethograms of locations and dyadic interactions. Location ethograms depict the time each mouse spent in the different ROIs of the SB, and dyadic interaction ethograms depict approaches, contacts, and chases between pairs in the group (Figure 2A-C). Dyadic interactions were modeled based on the relative ambulatory trajectories of the subjects, the distance ‘d’ between the mice, and the angle ‘θ’ of movement of each of the mice respective to each of the others (Figure 2B) (21). By definition, all dyadic interactions occur outside the ROIs ‘Large Nest’ and ‘Small Nest’, which are enclosed structures mice typically choose for rest. Visual inspection of the locations and dyadic interaction ethograms implies that EE groups exhibited lower exploratory activity and social interactions than SC controls (Figure 2D, E). Comparison between OxtR^-/-^ and OxtR^+/+^ groups (Figure 2F, G) revealed exaggerated exploratory activity and increased dyadic interactions in OxtR^-/-^ groups compared to OxtR^+/+^controls.

**Figure 2.**
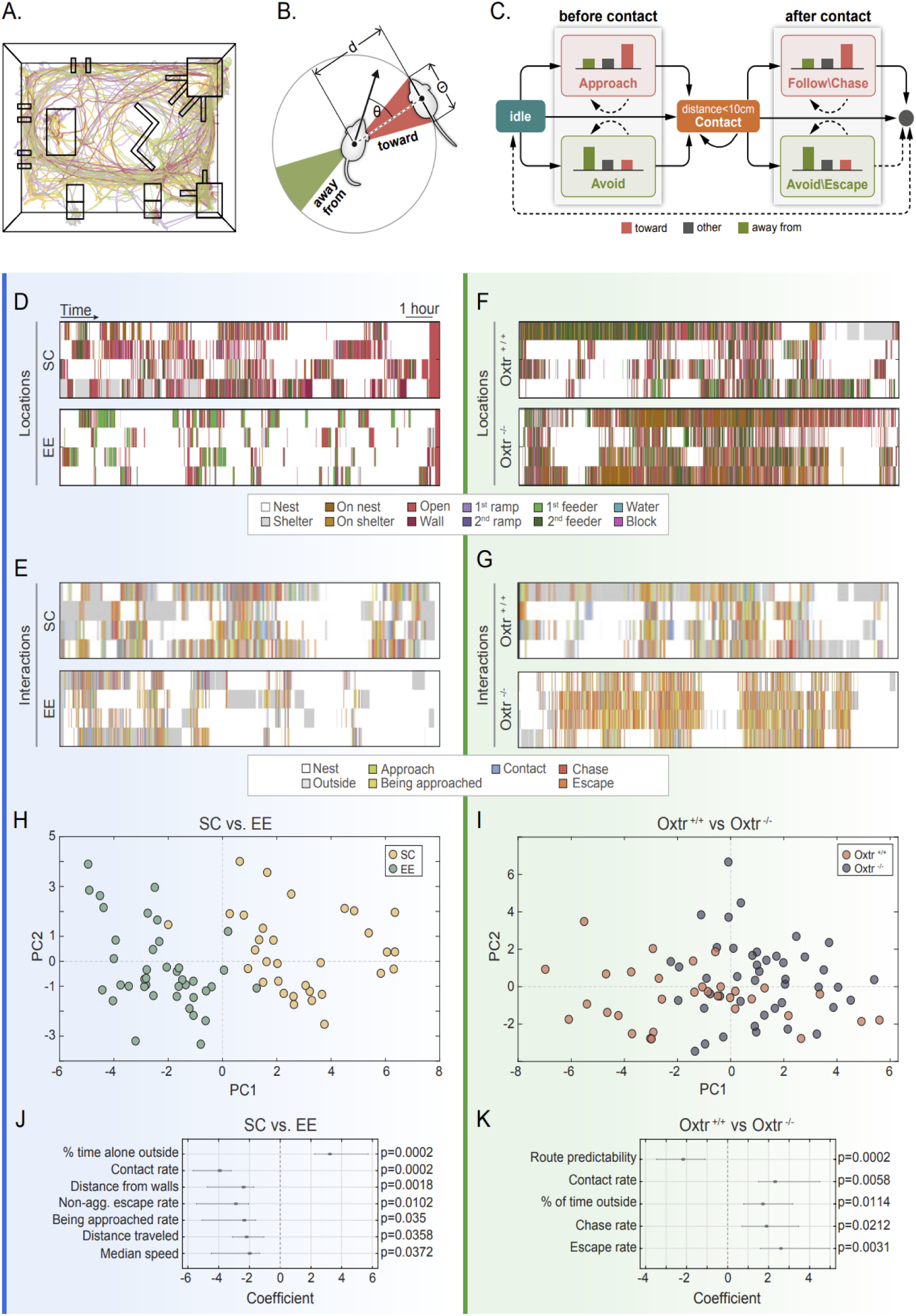
Exploratory and social activity is decreased following adolescence exposure to EE but increased in OxtR^-/-^. **(A)** A representative group depicting a 15-min segment of each mouse’s tracked paths (ambulatory trajectories). **(B, C)** Computational model of dyadic interactions. The algorithm used the distance (d) and the relative direction of the mouse respective to its conspecifics (θ) to determine if a mouse was moving toward another mouse, away from it, or neither. A contact is defined as a dyadic distance of less than 10 cm. A Hidden Markov Model (HMM) detects the events leading up to contact (Supplementary information). **(D)** Representative second-day EE and SC locations ethograms. **(E)** Representative second-day EE and SC dyadic interactions ethograms. **(F)** Representative second-day OxtR^-/-^ and OxtR^+/+^ locations ethograms. **(G)** Representative second-day OxtR^-/-^ and OxtR^+/+^ dyadic interactions ethograms. **(H, I)** A Scatterplot of the PCA’s PC1 and PC2 scores represents the behavioral readouts of EE vs. SC (H) and OxtR^-/-^ vs. OxtR^+/+^ (I). **(J, K)** Coefficients and 95% confidence intervals (CI) of logistic regression models that predict EE vs. SC (J) and OxtR^-/-^ vs. OxtR^+/+^ (K). Values correspond to behavioral readouts that, with the other behavioral readouts being equal, are more or less frequent in treatment groups (EE or OxtR^-/-^) compared to controls.

Next, we defined a set of 22 a priori chosen, automatically extracted behavioral readouts, and quantified the separation of those readouts in EE vs. SC and OxtR^-/-^ vs. OxtR^+/+^ groups in Euclidean space (19). These behavioral readouts included exploratory phenotypes and social interactions (*Supplementary Information*). Repeating this procedure multiple times with random partitioning of the data revealed that EE groups and OxtR^-/-^ groups show a statistically significant difference from their corresponding controls across the 22 dimensions of our dataset (Figure S1A; 10^7^ permutations; p=0.011, Figure S1B; 10^7^ permutations; 2.1e^-5^). In addition, to illustrate the separation in a lower dimensional space, we plotted the scores of the first two principal components (PC1 & PC2) emerging from a Principal Component Analysis (PCA) for each experiment and observed a considerable degree of separation in EE vs. SC (Figure 2H; Figure S2A; Figure S3A) and in OxtR^-/-^ vs. OxtR^+/+^ (Figure 2I; Figure S2B; Figure S3B), indicating the treatment effect (EE or OxtR^-/-^) constitutes a considerable source of variance in the dataset.

To identify the specific behavioral readouts that potentially drove the multivariate phenotypic differences, we fit a logistic regression model that attempts to predict the treatment group (EE vs. SC and OxtR^-/-^ vs. OxtR^+/+^) based on the set of behavioral readouts and estimates the contribution of each behavioral readout. Importantly, to remove potential collinearity between the behavioral features, which could potentially complicate the interpretation of the model coefficients, we first whitened the behavioral data using zero-phase component analysis (ZCA; see *Methods*). This model revealed that the most potent positive predictor variable for EE is the ‘fraction of time spent alone outside the nest’ (Figure 2J; β = 3.2, CI 95% 2.3-5.7, p=2×10^-4^). In other words, with all other behavioral variables held constant, EE mice tended to spend more time alone outside the nest. Readouts that were found to be significant negative contributors were ones that relate to social activity (’rate of contacts’, ‘being approached’ or ‘followed’) and exploratory behavior (’mean distance from walls’, ‘total distance traveled’, and ‘median speed’), (Figure 2J; rate of contacts: β = −3.9, CI 95% −5.7-(−3.2), p=2×10^-4^; being approached: β = −2.3, CI 95% −3.1 - (− 1.6), p=0.036; being followed: β = −2.9, CI 95% −5.4 - (−2.1), p=2×10^-4^). These results suggest that EE groups show social avoidance and reduced exploratory activity compared to SC.

In the OxtR^-/-^ experiment, the prediction of OxtR^-/-^ against OxtR^+/+^ brought up ‘rate of contacts’, ‘chasing’ and ‘escaping,’ as well as the ‘fraction of time spent outside the nest’ as significant positive predictor variables while ‘route predictability’ as a negative predictor variable (Figure 2K; rate of contacts: β = 2.3, CI 95% 1.5 - 4.5, p=0.006; chasing: β = 1.9, CI 95% 0.7 - 3.5, p=0.021; escaping: β = 2.6, CI 95% 1.6 - 5, p=0.031; fraction of time outside the nest: β = 1.7, CI 95% 0.79 - 3.2, p=0.011; route predictability: β = −2.2, CI 95% −3.5 - (−1.1), p=2×10^-4^). Together, these results suggest that, compared to the OxtR^+/+^ controls, OxtR^-/-^ groups show increased exploratory behavior and social activity, especially reciprocal aggressive chasing (Figure 2K). These findings suggest distinct patterns of group social dynamics in these two models of hyper-aggression that likely represent opposite behavioral effects. An overall reduction in exploration and social engagement in groups exposed to EE during adolescence and an increase in exploration and social engagement in OxtR^-/-^ groups.

### Divergent expression of hyper-aggression in EE and OxtR^-/-^ groups

Next, we directly compared the number of social encounters and aggressive chases in EE vs. SC and OxtR^-/-^ vs. OxtR^+/+^ groups. While the absolute rate of aggressive chasing in EE groups was drastically lower than in the SC controls (Figure 3A; permutation test, 1e7 permutations, p=0.0002), the EE mice engaged in significantly fewer contacts than mice in SC controls (Figure 3B; permutation test, 1e7 permutations, p=0.00002). Overall, the fraction of contacts that developed into an aggressive chase was significantly higher in EE groups compared to SC (Figure 3B inset; permutation test, 1e7 permutations, p=0.008). In contrast, the OxtR^-/-^ groups’ absolute rate of aggressive chasing was drastically higher than OxtR^+/+^ controls groups (Figure 3C; permutation test, 1e7 permutations, p=0.009) and they also engaged in significantly more contacts than OxtR^+/+^ controls groups (Figure 3D; permutation test, 1e7 permutations, p=0.037). However, the fraction of contacts that developed into chasing was not significantly different (Figure 3D inset; permutation test, 1e7 permutations, p=0.29). Hence, EE groups showed reduced social interactions and aggression compared to SC groups, with an increased likelihood of each interaction being aggressive. Such a strategy potentially minimizes the risks of injury. In contrast, compared to OxtR^+/+^ groups, the OxtR^-/-^ groups showed increased social interactions and aggression, putting individuals at a higher risk.

**Figure 3.**
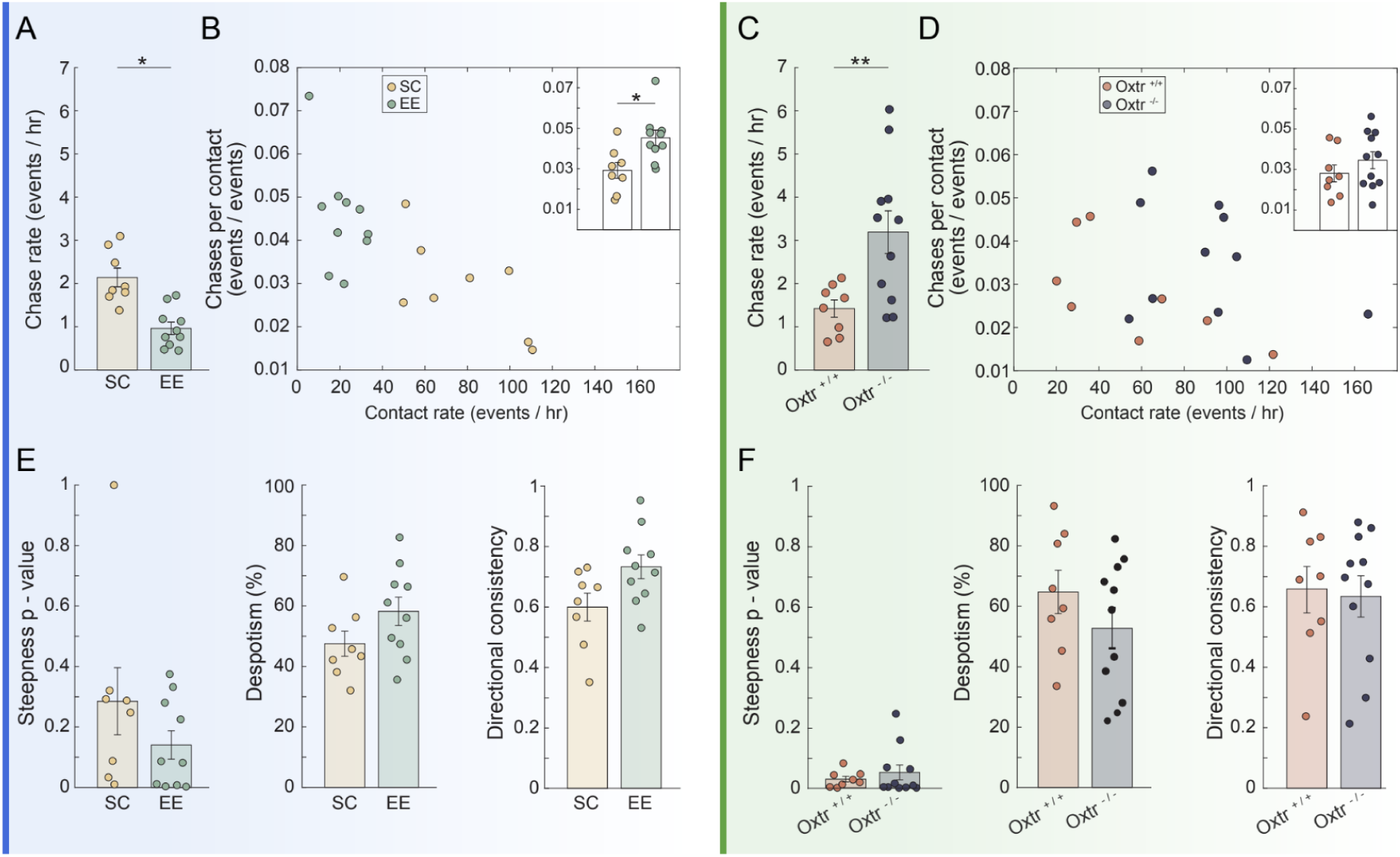
Divergent expression of hyper-aggression but enduring SDH in EE and OxtR^-/-^ groups. **(A-D) (A)** EE groups chase significantly less than SC groups. **(B)** While the rate of social contact is lower in EE compared to SC groups, the probability of a contact ending in aggression (a ‘chase’) is higher. The inset depicts a quantitative analysis of the chases per contact (by definition, each chasing event is preceded by a contact). **(C)** OxtR^-/-^ groups show an elevated rate of chases compared to OxtR^+/+^. **(D)** While the rate of contacts in OxtR^-/-^ is higher than that of OxtR^+/+^ groups, the probability of each contact ending in aggression (a ‘chase’) is similar. The inset depicts a quantitative analysis of the chases per contact. **(E)** No differences between EE and SC were found in the steepness of hierarchy, despotism, or directional consistency. **(F)** No differences were found between OxtR^-/-^ and OxtR in the steepness of hierarchy, despotism, or directional consistency. Values are mean ± s.e.m. * and ** differ from corresponding CTRL value (p< 0.05 and p<0.01 respectively)

### Social dominance hierarchies endure in both EE and OxtR^-/-^ groups

Next, we asked how social avoidance and increased aggression per contact in EE and increased social interactions and total levels of aggression in OxtR^-/-^ affect the social dominance hierarchies. Using ‘David’s Score’, an established method for measuring social dominance (28) and calculating the imbalance of weaning of aggressive chases in the SB, we previously demonstrated that groups of four male mice in the SB tend to form prominent and stable hierarchies (21). Here, based on David’s Score, we calculated a set of metrics reflecting different characteristics of the hierarchy in each group, namely, the steepness, despotism (the fraction of total encounters dominated by the top-ranking subject), and directional consistency (the fraction of encounters dominated by the higher ranked subject within each dyad) (Figure 3E, F). Using a randomization procedure, we calculated p-values for the steepness of EE vs. SC and OxtR^-/-^ vs. OxtR^+/+^ and found no significant difference (permutation test, 1e7 permutations, p= 0.22 and p=0.51 respectively). There were also no significant differences in despotism (p= 0.11 and p=0.0.24, respectively). Directional consistency was slightly higher in EE than SC (p=0.04) and did not differ between OxtR^-/-^ and OxtR^+/+^ groups (p=0.82).

### Temporal dynamic analysis of social dominance hierarchy

Although OxtR^-/-^ displays elevated social interactions, exploration, and aggression, they maintain SDH. This points to the importance of the hierarchical structure in the social dynamics of the group. SDH can be maintained with minimal aggression or continuous effort. Hence, we asked how efficient the maintenance of dominance hierarchy in EE vs. SC is and in OxtR^-/-^ vs. OxtR^+/+^. David’s Score for each group member is based on the total number of chases (‘wins’) and escapes (‘losses’) over the four days in the SB. We hypothesized that a temporal model of ‘wins’/’losses’ along the four days of the experiment could decipher intricate differences in hierarchy dynamics. We expected improved maintenance of dominance in EE and hence fewer temporal changes in EE compared to SC. Contrarily we expected more temporal changes in OxtR^-/-^ compared to OxtR^+/+^. To that aim, we analyzed the dyadic aggressive chases in the group using the Elo rating model. The Elo rating is a standard method for modeling SDH where the winner and loser scores are symmetrically adjusted according to the outcome and proportionally to the difference between the current ratings. As such, an unexpected outcome, such as a subordinate’s win against a powerful competitor, would result in a more significant adjustment, and vice versa (29). Visual qualitative inspection of the Elo rating trajectories of SC groups suggested unstable dynamics with frequent changes (Figure 4A). EE groups showed a divergence between one of the subjects and the rest, with minimal changes occurring from the second experimental day onwards (Figure 4B). By integrating the fluctuations of the Elo rating values throughout the experiment, we quantitatively estimated the inconsistency of each encounter outcome (i.e., the temporal instability of the hierarchy). As predicted, significantly fewer changes occurred in EE groups when compared to SC (Figure 4C, t(16)=-3.5, p=0.003). Qualitative (Figure 4D, E) and quantitative (Figure 4F) inspection of the OxtR^-/-^ compared to OxtR^+/+^ did not reveal apparent differences in OxtR^-/-^ compared to OxtR^+/+^ (t(17)=0.76, p=0.46).

**Figure 4.**
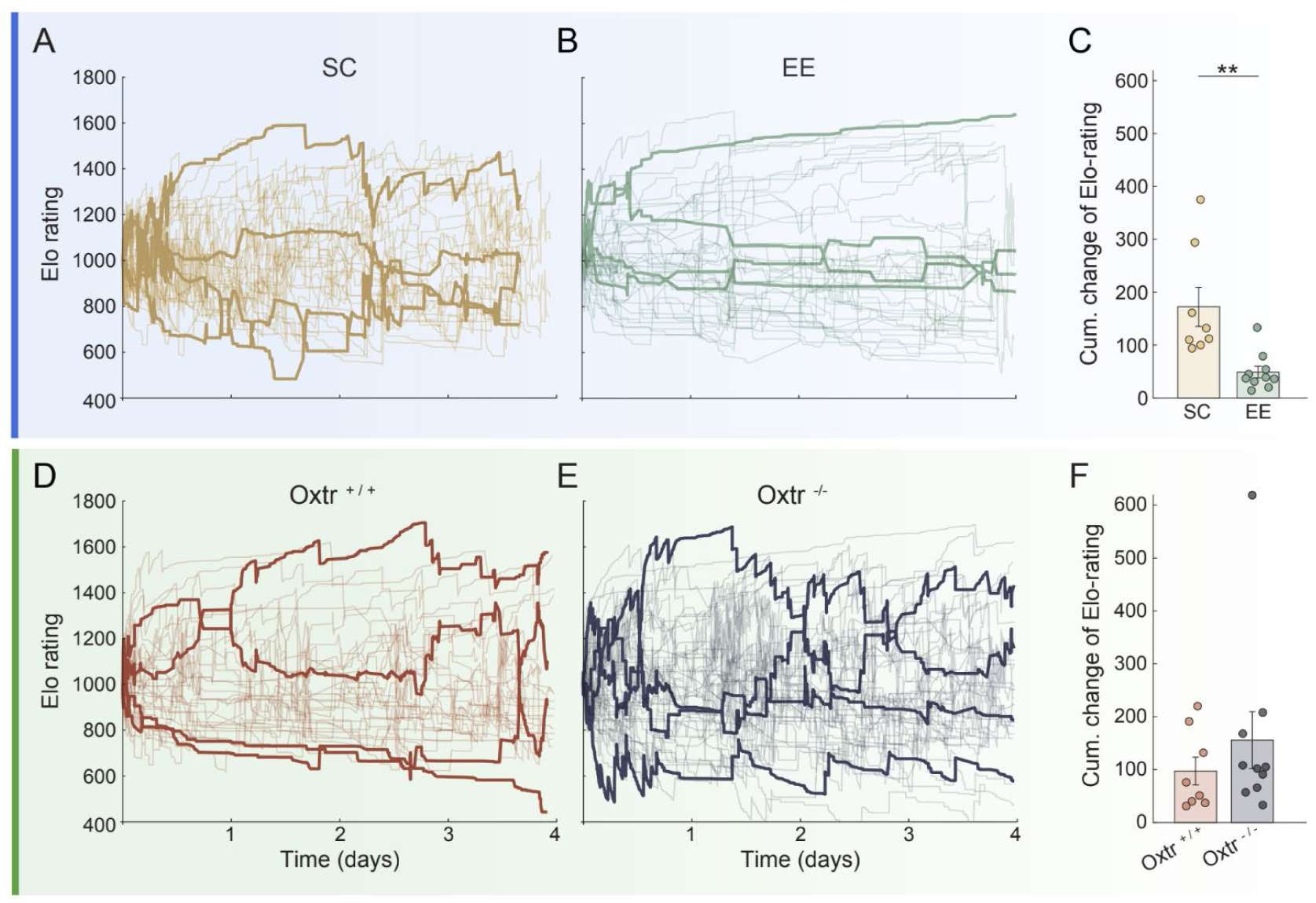
Temporal dynamic analysis of social dominance hierarchy. (**A, B**) Elo rating scores of all SC (A) and EE (B) mice. In bold, a single representative group of 4 mice. (**C**) Summation of the perturbations of the Elo rating values throughout the experiment in SC vs. EE. Fewer perturbations occurred in EE groups when compared to SC. (**D, E**) Elo rating scores of all OxtR^+/+^ (D) and OxtR^-/-^ (B) mice. In bold, a single representative group of 4 mice. (**F**) Summation of the perturbations of the Elo rating values throughout the experiment in OxtR^+/+^ vs. OxtR^-/-^. Values are mean ± s.e.m. * and ** differ from corresponding CTRL value (p< 0.05 and p<0.01, respectively)

### Differential access to resources is strengthened in EE compared to SC and weakened in OxtR^-/-^ compared to OxtR^+/+^ groups

SDH prioritizes access to food, water, territory, and mates, minimizing energy expenditure and the risk of injury through an aggressive encounter. Differential access to resources in the SB can be quantified as the time spent occupying the territory outside the shelter, being close to the feeders or water, and the patterns of ambulation trajectories outside the nest. To study the link between access to resources and dominance as measured using within-group chasing (David Score), we correlated (Pearson correlation) the DS with the within-group normalized values of the ‘percent of the time outside’ the ‘fraction of time near food or water’ and ‘route predictability’ (namely, the averaged mutual information of subsequent ROI transitions; see Supplementary information).

In mice exposed to early EE, the DS had a positive linear relationship with ‘time near the food and water’ and ‘fraction of time alone outside the nest’ and a negative relationship with ‘route predictability’. This indicates that dominants have more access to food and territory and that subordinates are restricted in their exploration. The correlations were more robust in EE than in SC (r_EE_= 0.63 vs. r_SC_= 0.27, r_EE_= 0.72 vs. r_SC_= 0.39, and r_EE_=-0.43 vs. r_SC_= −0.19 respectively), and the significance higher (p_EE_ = 1.29e-5 vs. p_SC_ = 0.129, p_EE_ =1.3e-7 vs. p_SC_ = 0.026 and p_EE_ =0.006 vs. p_SC_ = 0.3 respectively). This indicates that early exposure to EE strengthened Differential access to resources. However, the correlations were less robust in OxtR^-/-^ than in OxtR^+/+^ (r_OxtR+/+_=0.55 vs. r_OxtR-/-_=0.34, r_OxtR+/+_=0.61 vs. r_OxtR-/-_=0.11 and r_OxtR+/+_=-0.56 vs. r_OxtR-/-_=-0.05 respectively) and the p values were higher (p_OxtR+/+_ = 0.0009 vs. p_OxtR-/-_ = 0.02, p_OxtR+/+_ = 0.0002 vs. p_OxtR-/-_ = 0.46 and p_OxtR+/+_ = 0.0008 vs. p_OxtR-/-_ = 0.75 respectivly). This indicates that differential access to resources is weakened in OxtR^-/-^ compared to OxtR^+/+^ groups.

## Discussion

Though aggression is an efficient and common strategy in the everlasting struggle over limited resources, it must be tightly regulated to be displayed in the correct time and place (5,30). Disinhibition of aggression is a hallmark of many mental disorders, and the lack of appropriate behavioral mouse models hampers the effort to find an effective treatment for this symptom (4). Here we presented an ethologically relevant approach in mice that quantifies group behavior in a semi-natural setup and studies the reciprocal relationships between aggression, social avoidance, hierarchy, and access to resources. Such an approach enabled the study of the effect of EE and OxtR^-/-^ on adaptive aggression in a way that was not feasible before.

In the EE experiment, we exposed mice to a physically and socially enriched environment during adolescence. The EE paradigm can be considered a model for commensal habitats such as barns. Such habitats, typically rich in food and shelter, are characterized by intense competition between conspecifics (9). Exposure of mice to EE in laboratory conditions has been previously shown to enhance cognitive and physiological well-being, but also aggression and competition (11,27,31,32). Multivariate analysis of the behavioral data acquired in our SB paradigm revealed marked differences between EE and SC groups. Further investigation of specific readouts revealed that EE groups combine a social avoidance strategy with an increased probability of aggressive encounters once an encounter occurs (Figures 2 and 3). Such a strategy can strengthen the hierarchy and the differential access to resources while minimizing the risks of injuries (Figure 2; Figure S1). Social avoidance following intense aggression is a well-documented phenomenon. For instance, sequential exposures to intense aggression in the chronic social defeat paradigm can result in defeated mice avoiding unfamiliar males (33). Temporal analysis of winning/losing consistency in dyadic chases (Figure 4) confirmed that EE groups maintain hierarchy relationships more efficiently than SC groups and that Differential access to resources is strengthened in EE compared to SC (Figure 5A). These findings suggest that exposure to EE challenges during adolescence can reshape the group dynamics of adults to maintain differential access to resources with minimal but efficient levels of aggression.

**Figure 5.**
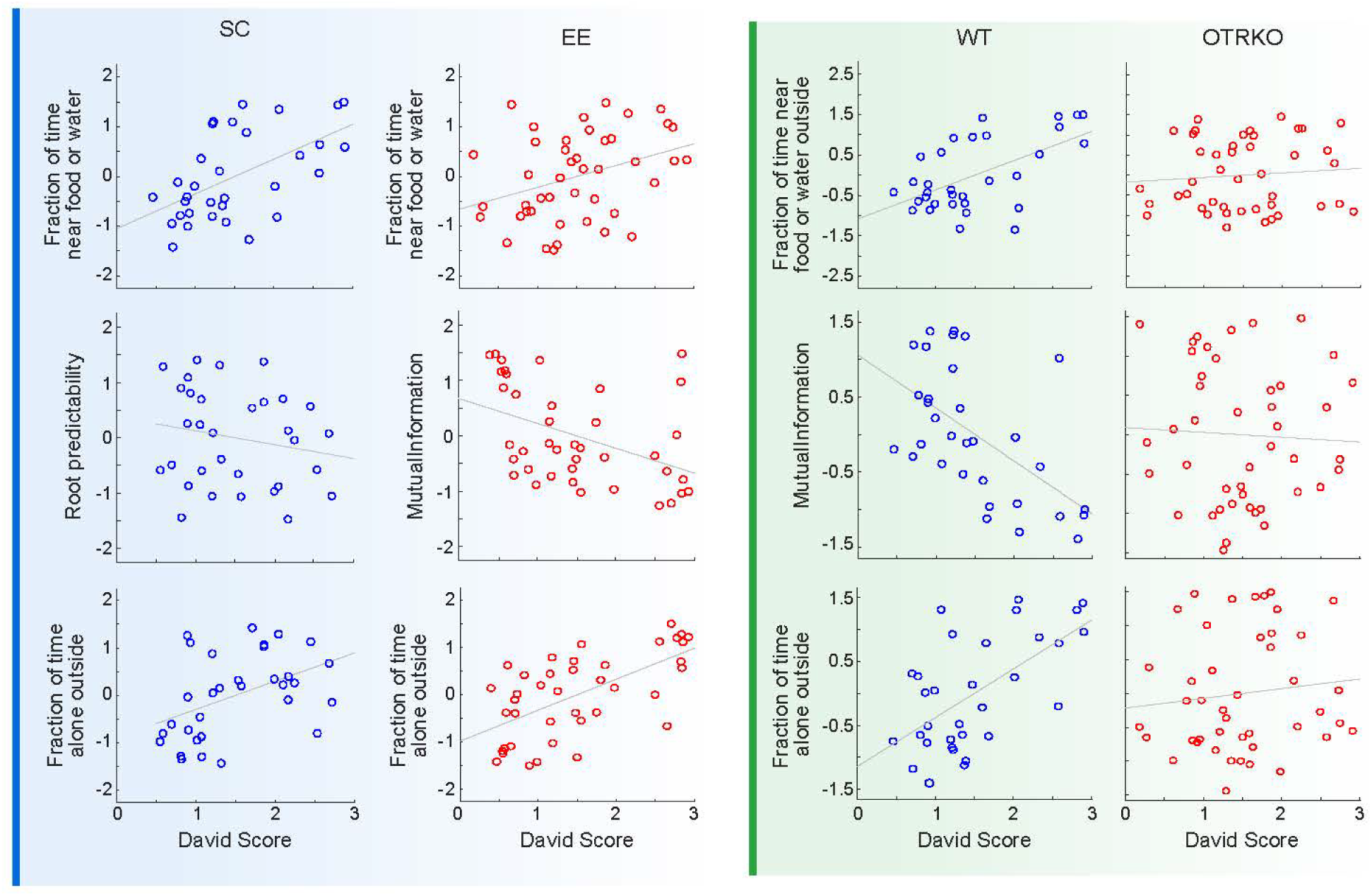
Organized priority order of access to resources in SC, EE, and OxtR^+/+^ groups but not in OxtR^-/-^ groups. **(A-C)** Scatter plots of time near food or water (A), exploration (B), and time outside (C) in SC and EE groups. **(D-F)** Scatter plots of time near food or water (D), exploration (E), and time outside (E) in OxtR^+/+^ and. OxtR^-/-^ groups.

In the second experiment, we modeled dysregulated aggression by introducing to the SB groups of mice that, since early development, lack the receptor for oxytocin (Figure 1). This receptor is widely expressed throughout the CNS, mediates a substantial part of the effects of oxytocin, and plays a pivotal role in many social behaviors (34–36). It is known that OxtR^-/-^ mice display excessive aggression in the RI paradigm (37), and it has also been suggested that the oxytocinergic system is involved in territorial behavior (24). Dimensionality reduction analysis of the multivariate behavioral data revealed that OxtR^-/-^ mice behave differently in the SB than the OxtR^+/+^ control groups. A logistic regression model to predict the treatment based on the behavioral readouts suggested that OxtR^-/-^ mice spend more time outside the nest, are more explorative, and are more aggressive to each other than OxtR^+/+^ groups (Figure 2 and Figure 3). Interestingly, OxtR^-/-^ groups can still maintain a hierarchy, suggesting that hierarchy is essential for group dynamics and survival even in genetic knockout mice lines that display severe behavioral dysfunction in classic tests (Figure 3). Nevertheless, Differential access to resources is strengthened in OxtR^-/-^ is weakened compared to OxtR^+/+^ mice. (Figure 5B). Overall, OxtR^-/-^ displays heightened aggression in a group context in the SB, and their hierarchy is not reflected by differential access to territory and food. These findings suggest that the increased aggression of OxtR^-/-^ mice is not constructive in terms of group coherence but rather a manifestation of irritation and anxiety that characterize this genetic manipulation. Of Note, the food supply was ad libitum, and the group members faced no outside danger like predation. It can be hypothesized that introducing food restriction and/or danger in Oxtr^-/-^ in the SB might induce differential access to resources or otherwise deteriorate the hierarchy.

In modern societies, aggression is strictly forbidden and tightly restricted by enforcing the rule of law. Nevertheless, violence is still a common symptom accompanying many mental illnesses, and its impact on the well-being of individuals and societies is tremendous. Since aggression, SDH, and competition over resources is common throughout the animal kingdom, it is reasonable to assume that the neuronal mechanisms underlying them are evolutionarily conserved to some extent between mammals. Hence, revealing the mechanisms that underlie ethologically-relevant dysregulated aggression in animal models is crucial for translational research. In mice, ‘escalated aggression’ towards an intruder in the RI test, such as excessive attacks, targeting vulnerable body parts, or ignoring submissive signals, has been suggested as a model of humans’ pathological aggression (1). However, severe aggression and even conspecific keeling are not rare among territorial species such as mice in their natural habitats (5). Therefore, a direct translation from what is considered human pathological aggression to animal models might not be adequate. Inhibitory mechanisms of aggression, such as SDH, can play a crucial survival role in humans as in any other territorial species. Hence a semi-natural approach that measures both aggression and SDH in groups of mice can be an essential complementary paradigm to reveal the intricate thresholds that separate adaptive from maladaptive aggression.

## Acknowledgments and Disclosures

A.C. is the incumbent of the Vera and John Schwartz Family Professorial Chair in Neurobiology at the Weizmann Institute of Science. This work was supported by Ruhman Family Laboratory for Research on the Neurobiology of Stress (to A.C.); research support from Bruno and Simone Licht; the Perlman Family Foundation, founded by Louis L. and Anita M. Perlman (to A.C.); the Adelis Foundation (to A.C.); and Sonia T. Marschak (to A.C.).

The authors report no biomedical financial interests or potential conflicts of interest.

## Article Information

## SUPPLEMENTARY INFORMATION

**Figure S1.**
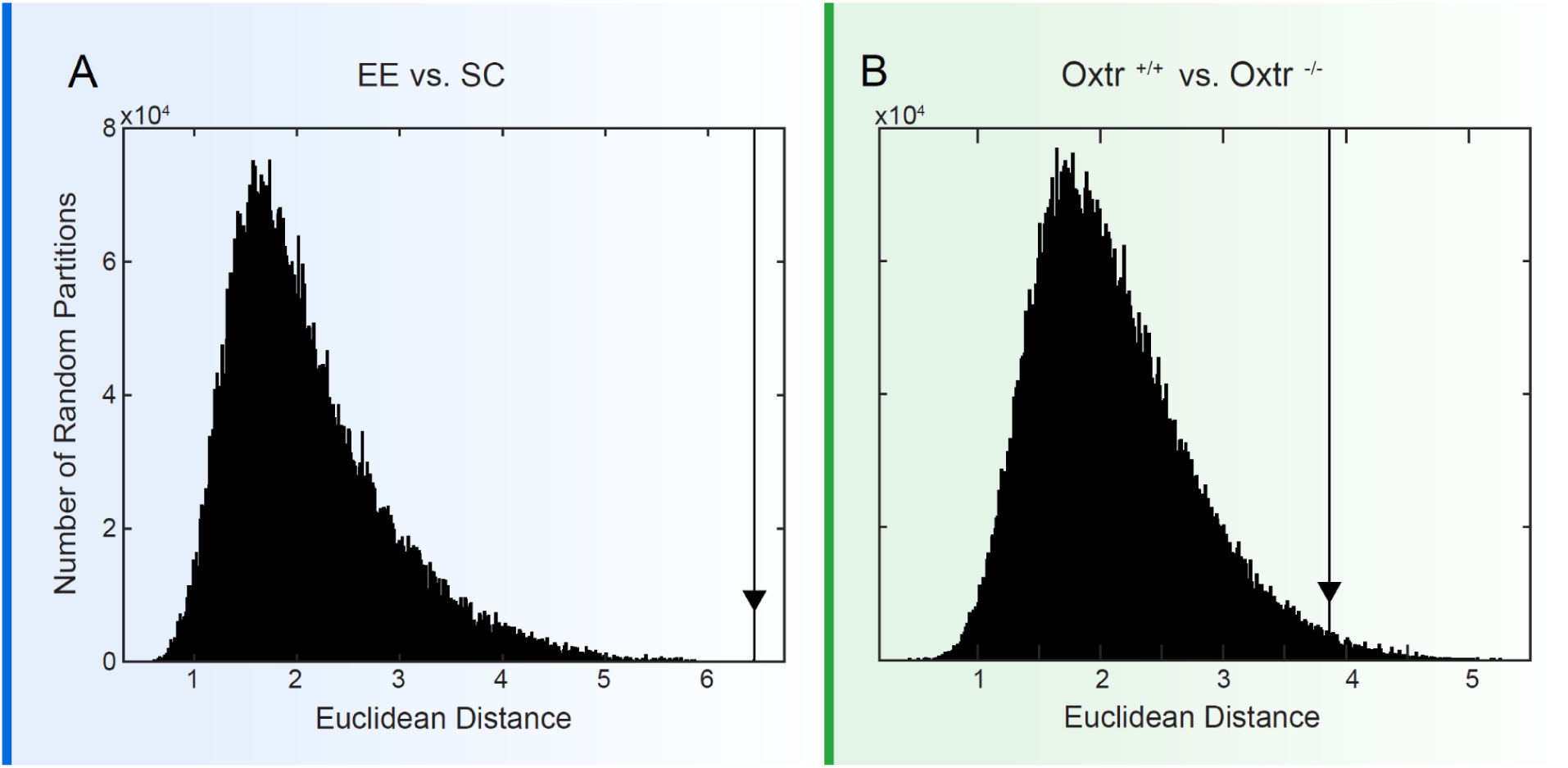
Euclidean distance between centroids of EE vs. SC and OxtR^+/+^ vs. OxtR^-/-^. Over the 21-dimensional space (number of behavioral readouts), EE was significantly different than SC (A), and OxtR^-/-^ was significantly different than OxtR^+/+^ (B). The Arrow line represents the distance corresponding to the actual data.

**Figure S2.**
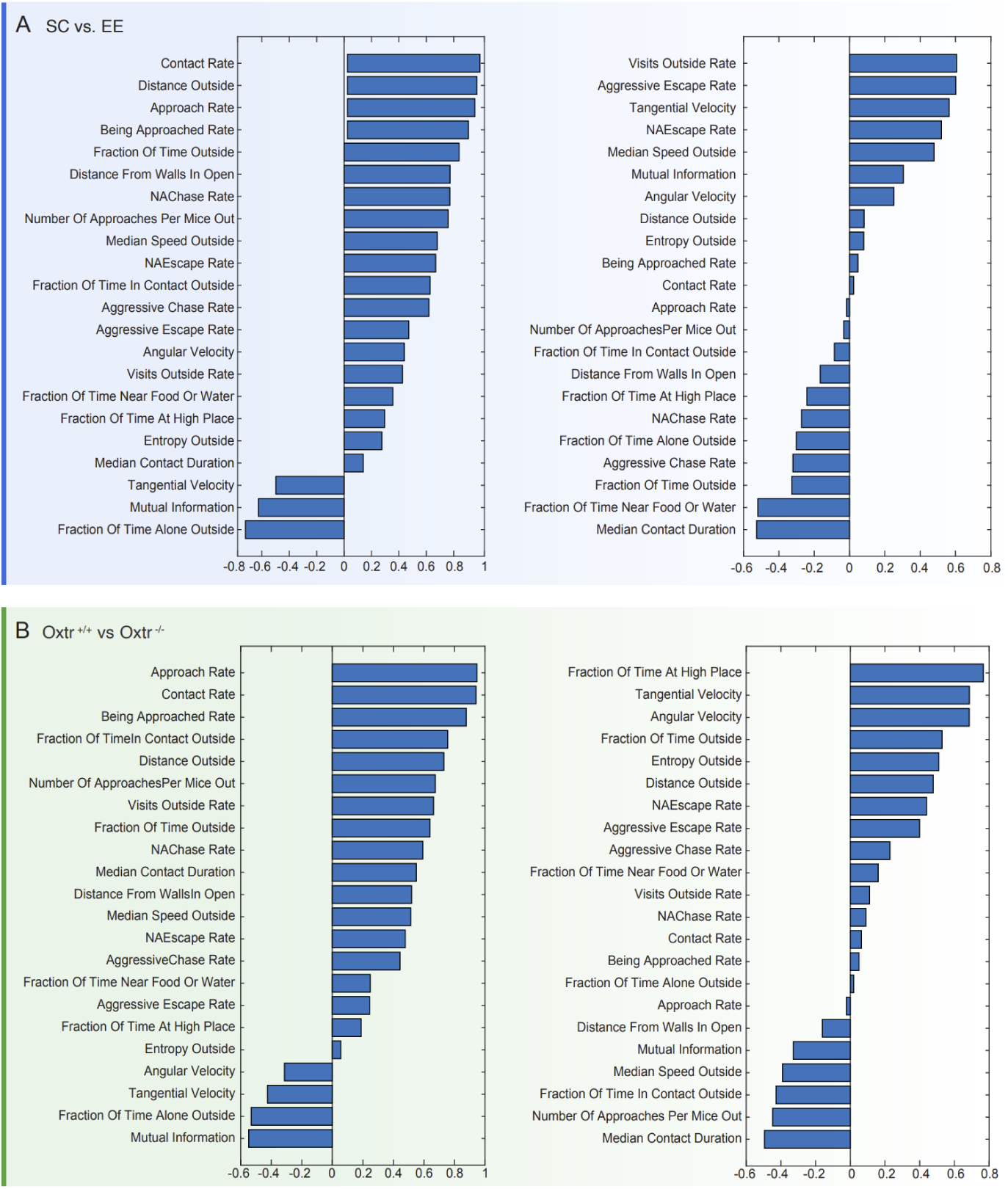
Graphical representation of the Pearson correlation coefficients with each of the 21 actual readouts with PC1 (left) and PC2 (right) scores.

**Figure S3.**
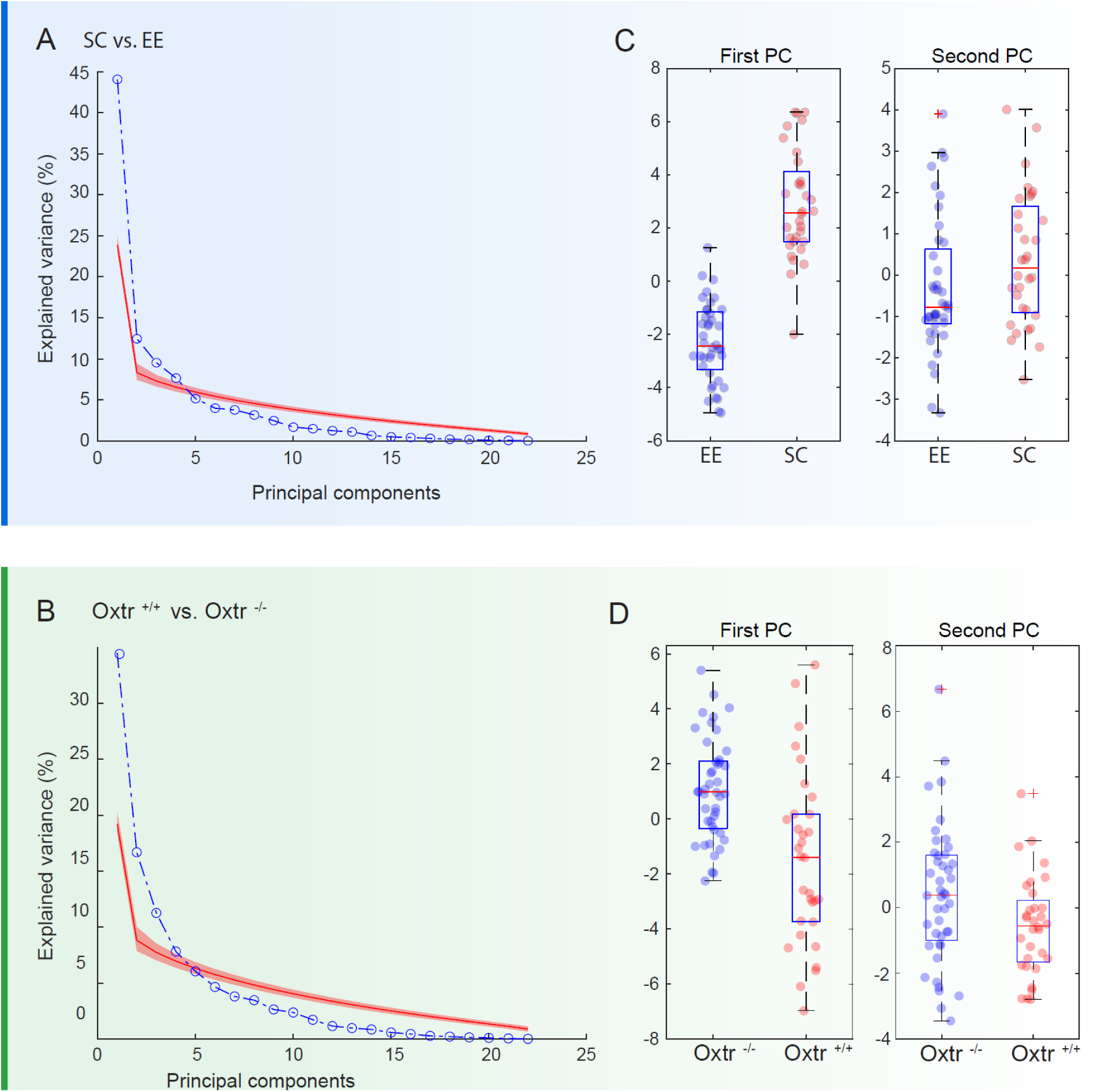
**(A,B)** Percentage of variance explained by each PC given by PCA performed on actual, as compared to permuted, data. Values of permuted data (red) represent means and 95% confidence interval. in EE vs. SC (A) and in OxtR^+/+^ vs. OxtR^-/-^ (B). (**C,D**) Box plots depicting the projected scores on each PC, separately. EE and SC (C), OxtR^+/+^ vs. OxtR^-/-^ (D).

## SUPPLEMENTARY MATERIALS and METHODS

### Animals

All experimental protocols were approved by the Institutional Animal Care and Use Committee of The Weizmann Institute of Science.

EE experiment: Adult male ICR mice (Harlan Laboratories, Jerusalem, Israel) were used. Throughout the experiments, the animals were maintained in a temperature-controlled mouse facility (22°C ± 1) on a reverse 12 hr light–dark cycle. Food and water were given ad libitum.

OxtR^-/-^ experiment: OxtR^-/-^ mice, which were originally generated by Lee et al. (2008), carried the mutation on the genetic background of the 129/Sv line and were crossed with ICR mice (Harlan Laboratories, Jerusalem, Israel) to get heterozygous on a mixed background (F1). Experimental mice were produced by crosses of heterozygous F1 parents. To form experimental groups of four male mice, two pairs of littermates obtained from a separate family were weaned into the same cages. Since weaning and before the experiments, the mice were housed in groups. Following color marking, OxtR^-/-^ mice were transferred from the regular to a reverse light cycle (10^00^-22^00^ dark, 22^00^-10^00^ light).

### Environmental enrichment cages

At the age of weaning (three wks), mice were randomly distributed into two types of groups: standard conditions (SC) mice that were housed in groups of four in standard laboratory cages and enriched environment (EE) mice that were housed in groups of 16 male mice in a relatively spacious and complex cage, with a variety of objects such as shelters, tunnels, running wheels, and mouse nest boxes (Sztainberg and Chen 2010). After six weeks, EE mice were randomly divided into groups of four, color marked, and introduced to the social box.

### The social box

#### Color markings of the fur

Mice were mildly anesthetized with isoflurane. Their eyes were protected against drying using eye gel (viscotears liquid gel; Alcon). The fur of the mice was stained using a regular brush with fluorescent semi-permanent hair dyes that glow under black light. The fur was dried with a fan (low power and heat) for three min. After awakening, mice were kept in separate carton boxes for one hr before the reunion. Mice were introduced to the arena 1 week after the fur staining. The dyes (composed of natural ingredients) were “Electric banana” (HCR 11012), Atomic™ Turquoise (AMPLIFIED™ HCR11002), Ultra™ Violet (AMPLIFIED™ ACR-71031), and Pillarbox™ Red (HCR 11020) from Tish & Snooky’s (https://manicpanic.com; New York NY, USA). Mice were identified and tracked automatically according to their fur colors, learned from labeled data.

#### The arenas

The SB consisted of an open 70 cm × 50 cm × 50 cm box and included the following objects: A z-shaped wall, a water dispenser, two feeders, a small nest and a large nest, an elevated block, and two elevated ramps. Food and water were given ad libitum. During the 12h dark phase, UVA fluorescent lamps (18 W) were placed three meters above the arena’s floor to illuminate the surrounding area with 370–380 nm ‘black light’. During the 12 hours light phase, the arena was illuminated with two regular (18W) fluorescents. A black curtain was drawn from the fluorescent lamps down to the arena to avoid reflections from white objects in the room. A color-sensitive camera (Panasonic Color CCTV, WV-CL924AE) was placed 1 m above the arena. The camera analog input was converted to digital information with a digitizer (Picolo Diligent frame grabber board) and recorded on a standard computer.

#### Mouse tracking

Mouse trajectories were automatically detected offline using custom software in MATLAB (R2019a, MathWorks, Natick, MA, USA). The mice were identified according to their fur colors, learned from labeled data. Because of the low signal-to-noise ratio, due to the dim lighting and the camera’s sensitivity, some frames had reflection artifacts or missing parts. To overcome this noise, we used a Bayesian model to infer the most likely location of a mouse given the observed location of connected colored blobs (see Shemesh et al. e Life).

### Identification and classification of behavior

We defined a set of 22 a priori chosen behavioral readouts as described in Anpilov 2019. The behavioral readouts included both Individual automatic behavioral readouts as well as pairwise automatic behavioral readouts.

#### Individual automatic behavioral readouts

##### Time outside

Fraction of time that the mouse spends outside the nest (% of total). Rate of visits outside: The rate at which the mouse exits the nest (1/hour).

##### ROI exploration

Quantifies the amount of exploration the mouse is doing. Measured as the entropy of the probability of being in each of the 11 regions of interest (ROIs) (excluding the nest). Mice that spend the same amount of time in all regions will get the highest score, while mice that spend all their time in a single ROI will be scored zero. When normalized to the time outside, the entropy computation also differed by ignoring the probability of being inside the nest (bits/hour). Distance: The total distance traveled by the mouse while outside the nest. The mice locations were sampled once every second (m) to smooth the tracking.

##### Median speed

Median speed while outside the nest. To smooth the computation of the speed, the mice locations were sampled once every second (m/sec).

##### Tangential velocity

The tangential component of the speed, or the part of speed perpendicular to the previous direction of movement (m/sec).

##### Angular velocity

The rate of change in the direction of the mouse (rad/sec). Food or water: Time spent next to the feeders or water bottles (au).

##### Elevated area

Time spent on an elevated object in the arena: ramps or block (au). Distance from walls: Average distance from the walls while in the open area (cm).

#### Pairwise automatic behavioral readouts

We automatically identified and classified interactions between mice as events in which the distance between a pair of mice (d) was shorter than 10 cm. We then used the movement direction of one mouse relative to another mouse to identify the nature of the contact for either of the mice. If, for mouse A, the projection of the direction of its movement relative to mouse B was small enough, then it was considered as moving towards B; if it was moving away from it; otherwise it was assumed idle with respect to the other mouse (and were found by optimization).

As described by Forkosh et al. (21), to classify aggressive and non-aggressive contacts, we first used a hidden Markov model to identify post-contact behaviors in which mouse A was moving towards B, and B was moving away from A (i.e., A was following B). We then used 500 manually labeled events to learn aggressive and non-aggressive post-contact behavior statistical classifiers. For each event, we estimated a range of parameters, including individual and relative speed, distance, etc. We optimized a quadratic discriminant classifier, a k-nearest neighbor algorithm based on these parameters, and a decision-tree classifier that used these parameters at each tree intersection. We found that for a test set of 1000 events, none of these classifiers were accurate enough individually, but that a combined approach in which we labeled an event as ‘aggressive’ if any of the classifiers labeled it as such – gave ∼80% detection with 0.5% false alarms.

##### Contact rate

Number of contacts the mouse had. A contact is defined as conspecifics being less than 10 cm apart while both are outside the nest (1/hour).

##### Time in contact

Fraction of time a mouse is in contact with other mice outside the nest (1/hour). Median contact duration: Median duration of contacts. The contact duration does not include when the mouse approached, moved away from, or chased a conspecific (seconds).

##### Chase rate

Chases are interactions that end with the mouse aggressively pursuing a conspecific. Aggressiveness was determined using a classifier trained on 500 manually scored samples (1/hour).

##### Escape rate

Number of times the mouse was aggressively chased by a conspecific (1/hour). Chase rate (non-aggressive): Number of times the mouse has chased a conspecific at the end of a contact in a non-aggressive way, as determined by the classifier (1/hour).

##### Escape rate (non-aggressive)

Number of times the mouse was chased by a conspecific at the end of a contact in a non-aggressive way, as determined by the classifier (1/hour).

##### Approach rate

An approach is a directed movement of the mouse towards a conspecific that ends in contact. Not all interactions necessarily start with an approach, while others might start mutually with conspecifics approaching each other (1/hour).

##### Approach rate (per mice out)

number of approach events normalized by time outside the nest with at least one more conspecific (1/hour).

##### Rate of being approached

Number of times the mouse was approached by a conspecific (1/hour). Alone outside: Fraction of time the mouse is outside the nest while all other mice are in the nest (au).

### Quantification and statistical analysis

Multi-variate separation: Was calculated as Euclidean distance between the centroids of Cre^+^ and Cre^-^ mice in 21 dimensions of the automatically collected behavioral readouts. Statistical inference was based on comparison to a null distribution of Euclidean distances corresponding to 1×10^5^ randomly reshuffled partitions.

Principal component analysis (PCA): principal component analysis (PCA) was performed on the raw collected data from the above-listed 21 behavioral readouts. on the Z-scores. The explained variance was independently calculated from one million randomly reshuffled versions of the original dataset to determine the percentage of variance explained by each principal component (PC) in a random dataset. Specifically, reshuffled datasets were created by independently reshuffling the variables within each observation.

Elo rating: The measurement of hierarchy instability was based on outcomes of aggressive dyadic interactions, namely ‘chases’ or ‘escapes’. For each dyad, we tested the assumption that both mice have equal probabilities of chasing its counterpart. When the assumption was met, the mice were assigned similar ranks. However, upon a disproof of this assumption, the mice were divided into different ranks, with the mouse that chased more receiving a higher rank. This was repeated for every day of the experiment. The numbers of intra-dyadic rank change between subsequent days were summed throughout the experiment. For each group, the resulting values were divided by the total number of possible dyads to receive a proportion of dyads that changed during the experiment, thus representing the instability of the dominance hierarchy.

By treating chasing and escaping as ‘wins’ and ‘losses’, this principle can be quantitatively represented by the Elo rating, a method used to assign rating scores in chess competitions. Briefly, every participant is assigned the same initial rating. Upon each subsequent encounter, the ratings of the winner and the loser are symmetrically adjusted according to the outcome and proportionally to the difference between the current ratings.

#### David score calculation

David’s score (DS) was calculated based on aggressive chases, as described in de Vries et al. (28). First, the dyadic proportions of chases are calculated. The proportion of chases by the mouse *i* in its interactions with another’s mouse *j* (*Pij*) is the number of chases of *i* after *j* divided by the total number of interactions between *i* and *j* (*n_ij_*). For each group member, DS is calculated with the following formula: *DS*=*w*2+*w*−*l*−*l*2. Here, *w* represents the sum of *Pij* values: *w*=Σ(*j*=1..*n*;*j*≠*i*). *w*2 is the sum of *Pij* values weighted by *w* values of the interactants: *w*2=Σ(*j*=1..*n*;*j*≠*i*). Calculations of *l* and *l*2 are similar to *w* and *w*2, but for the number of escapes instead of the number of chases. Further corrections for the number of interactions and subjects were done as proposed in (28).

The steepness of the social hierarchy: was characterized as described in de Vries et al., (28) by using the slope of a line fitted to the DS from a ranked DS using Ordinary Least Squares regression. We implemented this procedure made available in the open-source ‘steepness’ R package (Leiva and de Vries, 2014), whose output ranges between 0 and 1, with one meaning a very steep hierarchy in which power is unequally distributed between dominant and subordinate individuals. Despotism: Was defined as the fraction of the group’s total number of chases initiated by the highest-ranking individual. Its values also range between 0 and 1, in which one represents the presence of a powerful alpha who initiates all chases.

Directional consistency: Was calculated using the average fraction of pairwise social interactions that occur in the direction from the individual who displayed more agonistic behavior to the individual who displayed fewer instances (van Hooff and Wensing, 1987; Williamson et al., 2016). A directional consistency equal to one indicates that all agonist interactions are directed from an individual with a higher DS to one with a lower DS. We calculated directional consistency using functions made available in the R package ‘compete’ (Curley et al., 2015).

### Logistic and linear regression models

To quantify the relationship between the group types (EE vs. SC and OxtR^-/-^ vs. OxtR^+/+^) and the behavioral readouts measured, logistic regression models were fit with group type as the label and behavioral readouts as the features, using the Matlab (2021b) function ‘fitclinear’ with Lasso regularization and the default parameters.

Similarly, ‘fitrlinear’ was used to fit linear regression models with social dominance (David’s scores) as the label and occupation of crucial resources as the features.

Leave-one-out cross-validation (CV) was used to estimate the performance of these models. Next, the value of the regularization parameter was chosen based on CV performance to avoid overfitting and assert that only truly predictive features will have significant coefficients, and the coefficients of these well-regularized models and their 95% confidence intervals were then used to examine the relationship between the variables. Importantly, to remove potential colinearity between the behavioral readouts, which could potentially complicate the interpretation of the model coefficients, the behavioral data were first whitened using zero-phase component analysis (ZCA), and the models were fit to these whitened data.

For inferential comparisons between two samples, the Monte Carlo permutation test of means (1×10^7^ permutations) was carried out. All the above analyses were performed using MATLAB (R2019a, MathWorks, Natick, MA, USA).

## References

1. Miczek KA, De Boer SF, Haller J (2013): Excessive aggression as model of violence: A critical evaluation of current preclinical methods. Psychopharmacology (Berl) 226: 445–458.

2. Ramirez JM (2009): Some dychotomous classifications of aggression according to its function. Journal of Organisational Transformation & Social Change 6: 85–101.

3. Allen JJ, Anderson CA (2017): Aggression and Violence: Definitions and Distinctions. The Wiley Handbook of Violence and Aggression. John Wiley & Sons, Ltd, pp 1–14.

4. Siever LJ (2008, April): Neurobiology of aggression and violence. American Journal of Psychiatry, vol. 165. pp 429–442.

5. Gómez JM, Verdú M, González-Megías A, Méndez M (2016): The phylogenetic roots of human lethal violence. Nature 538: 233–237.

6. Kwiatkowski CC, Akaeze H, Ndlebe I, Goodwin N, Eagle AL, Moon K, et al. (2021): Quantitative standardization of resident mouse behavior for studies of aggression and social defeat. Neuropsychopharmacology 46: 1584–1593.

7. Lischinsky JE, Lin D (2020): Neural mechanisms of aggression across species. Nat Neurosci 23: 1317–1328.

8. Kondrakiewicz K, Kostecki M, Szadzińska W, Knapska E (2019): Ecological validity of social interaction tests in rats and mice. Genes Brain Behav 18: 1–14.

9. Bronson FH (1979): The Reproductive Ecology of the House Mouse. QUARTERLY REVIEW OF BIOLOGY, vol. 54.

10. Howerton CL, Garner JP, Mench JA (2008): Effects of a running wheel-igloo enrichment on aggression, hierarchy linearity, and stereotypy in group-housed male CD-1 (ICR) mice. Appl Anim Behav Sci 115: 90–103.

11. Van Loo PLP, Mol JA, Koolhaas JM, Van Zutphen BFM, Baumans V (n.d.): Modulation of Aggression in Male Mice: Influence of Group Size and Cage Size.

12. McQuaid RJ, Dunn R, Jacobson-Pick S, Anisman H, Audet MC (2018): Post-weaning environmental enrichment in male CD-1 mice: Impact on social behaviors, corticosterone levels and prefrontal cytokine expression in adulthood. Front Behav Neurosci 12.

13. Dwortz MF, Curley JP, Tye KM, Padilla-Coreano N (2022): Neural systems that facilitate the representation of social rank. Philosophical Transactions of the Royal Society B: Biological Sciences, vol. 377. Royal Society Publishing.

14. Caroline Blanchard D, Blanchard RJ, BLANCHARD Behavioral RJ (n.d.): Behavioral Correlates of Chronic Dominance-Subordination Relationships of Male Rats in a Seminatural Situation. Neuroscience & Biobehavioral Reviews, vol. 14.

15. Fan Z, Zhu H, Zhou T, Wang S, Wu Y, Hu H (2019): Using the tube test to measure social hierarchy in mice. Nat Protoc 14: 819–831.

16. Shemesh Y, Chen A (2023): A paradigm shift in translational psychiatry through rodent neuroethology. Molecular Psychiatry, 1–11.

17. Shemesh Y, Sztainberg Y, Forkosh O, Shlapobersky T, Chen A, Schneidman E (2013): High-order social interactions in groups of mice. Elife 2013: 1–19.

18. Shemesh Y, Forkosh O, Mahn M, Anpilov S, Sztainberg Y, Manashirov S, et al. (2016): Ucn3 and CRF-R2 in the medial amygdala regulate complex social dynamics. Nat Neurosci 19: 1489– 1496.

19. Anpilov S, Shemesh Y, Eren N, Harony-Nicolas H, Benjamin A, Dine J, et al. (2020): Wireless Optogenetic Stimulation of Oxytocin Neurons in a Semi-natural Setup Dynamically Elevates Both Pro-social and Agonistic Behaviors. Neuron 107: 644–655.e7.

20. Karamihalev S, Brivio E, Flachskamm C, Stoffel R, Schmidt M V., Chen A (2020): Social dominance mediates behavioral adaptation to chronic stress in a sex-specific manner. Elife 9: 1– 18.

21. Forkosh O, Karamihalev S, Roeh S, Alon U, Anpilov S, Touma C, et al. (2019): Identity domains capture individual differences from across the behavioral repertoire. Nat Neurosci 22: 2023– 2028.

22. Ne’eman R, Perach-Barzilay N, Fischer-Shofty M, Atias A, Shamay-Tsoory SG (2016): Intranasal administration of oxytocin increases human aggressive behavior. Horm Behav 80: 125–131.

23. Lee HJ, Caldwell HK, Macbeth AH, Tolu SG, Young WS (2008): A conditional knockout mouse line of the oxytocin receptor. Endocrinology 149: 3256–3263.

24. Wirth S, Soumier A, Eliava M, Derdikman D, Wagner S, Grinevich V, Sirigu A (2021): Territorial blueprint in the hippocampal system. Trends Cogn Sci 1–12.

25. Jurek B, Neumann ID (2018): THE OXYTOCIN RECEPTOR: FROM INTRACELLULAR SIGNALING TO BEHAVIOR. Physiol Rev 98: 1805–1908.

26. Samuni L, Preis A, Mundry R, Deschner T, Crockford C, Wittig RM (2017): Oxytocin reactivity during intergroup conflict in wild chimpanzees. Proc Natl Acad Sci U S A 114: 268–273.

27. Sztainberg Y, Chen A (2010, September): An environmental enrichment model for mice. Nature Protocols, vol. 5. pp 1535–1539.

28. De Vries H, Stevens JMG, Vervaecke H (2006): Measuring and testing the steepness of dominance hierarchies. Anim Behav 71: 585–592.

29. Albers PCH, De Vries H (2001): Elo-rating as a tool in the sequential estimation of dominance strengths. Animal Behaviour, vol. 61. Academic Press, pp 489–495.

30. Wang F, Kessels HW, Hu H (2014, November 1): The mouse that roared: Neural mechanisms of social hierarchy. Trends in Neurosciences, vol. 37. Elsevier Ltd, pp 674–682.

31. Ratuski AS, Weary DM (2022, February 1): Environmental Enrichment for Rats and Mice Housed in Laboratories: A Metareview. Animals, vol. 12. MDPI.

32. Dubois F, Giraldeau LA (2005): Fighting for resources: The economics of defense and appropriation. Ecology 86: 3–11.

33. Golden SA, Covington HE, Berton O, Russo SJ (2011): A standardized protocol for repeated social defeat stress in mice. Nat Protoc 6: 1183–1191.

34. Guzmán YF, Tronson NC, Jovasevic V, Sato K, Guedea AL, Mizukami H, et al. (2013): Fear-enhancing effects of septal oxytocin receptors. Nat Neurosci 16: 1185–1187.

35. Grinevich V, Neumann ID (2021, January 1): Brain oxytocin: how puzzle stones from animal studies translate into psychiatry. Molecular Psychiatry, vol. 26. Springer Nature, pp 265–279.

36. Kosfeld M, Heinrichs M, Zak PJ, Fischbacher U, Fehr E (2005): Oxytocin increases trust in humans. Nature 435: 673–676.

37. Takayanagi Y, Yoshida M, Bielsky IF, Ross HE, Kawamata M, Onaka T, et al. (2005): Pervasive Social Deficits, but Normal Parturition, in Oxytocin Receptor-Deficient Mice. Retrieved from www.pnas.orgcgidoi10.1073pnas.0505312102

